# Prophage elements function as reservoir for antibiotic resistance and virulence genes in nosocomial pathogens

**DOI:** 10.1101/2020.11.24.397166

**Authors:** Kohei Kondo, Mitsuoki Kawano, Motoyuki Sugai

## Abstract

Prophages are often involved in host survival strategies and contribute toward increasing the genetic diversity of the host genome. Prophages also drive horizontal propagation of various genes as vehicles. However, there are few retrospective studies contributing to the propagation of antimicrobial resistance (AMR) and virulence factor (VF) genes by prophage. In this study, we extracted complete genome sequences of seven pathogens, including ESKAPE bacteria and *Escherichia coli* deposited in a public database, and examined the distribution of both AMR and VF genes in certain genomic regions of prophage, including prophage-like element. We found that the ratios of AMR and VF genes greatly varied among the seven species. More than 55% of *Enterobacter cloacae* strains had VF genes, but only 0.8% of *Klebsiella pneumoniae* strains had VF genes from prophages. The prophage types carrying AMR genes were detected in a broad range of hosts, whereas prophages containing VF genes were conserved in only one or two species, suggesting that distribution patterns of prophages were different between prophages encoding AMR or VF genes. We also found that the prophage containing class 1 integrase possessed a significantly higher number of AMR genes than prophages with no class 1 integrase. Moreover, AMR genes in the prophage were located near transposase and integrase. The results of this study reveal a comprehensive picture of AMR and VF genes present in prophage elements and provide new insights into the horizontal transfer of genes associated with antimicrobial resistance and pathogenicity.

**Importance:** Although we believe phages play an important role in horizontal gene transfer in exchanging genetic material, we do not know the distribution of the antimicrobial resistance and/or virulence genes in prophages. We collected different prophage elements from the complete genome sequence of seven species – *Enterococcus faecium, Staphylococcus aureus, Klebsiella pneumoniae, Acinetobacter baumannii, Pseudomonas aeruginosa*, and *Enterobacter cloacae*, as well as *Escherichia coli* –, and characterized the distribution of antimicrobial resistance and virulence genes encoded in the prophage region. While virulence genes in prophage were found to be species-specific, antimicrobial resistance genes in prophages were highly conserved in various species. Integron structure was detected within prophage regions in almost all of the genera. Maximum of 11 antimicrobial resistance genes were found in a single prophage region, suggesting that prophages act as a reservoir for antimicrobial resistance genes. Our results highlight new insights on prophages as horizontal gene carriers.

## Introduction

Antimicrobial resistance (AMR) is a global public health issue. In recent years, ESKAPE pathogens (*Enterococcus faecium, Staphylococcus aureus, Klebsiella pneumoniae, Acinetobacter baumannii, Pseudomonas aeruginosa*, and *Enterobacter* spp.) have become a threat since they are the leading cause of nosocomial infection and easily escape from authentic chemotherapy due to their antimicrobial-resistant phenotype and many countries have faced difficulties in controlling these pathogens (1, 2). In Japan, *Escherichia coli* has replaced *Staphylococcus aureus* as the primary pathogen isolated from clinical samples in hospitals since 2018, and isolation of third-generation cephalosporin-or quinolone-resistant *E. coli* continues to increase in Japan (https://janis.mhlw.go.jp). Moreover, the extended-spectrum of ß-lactamase-producing *E. coli* is spreading worldwide (3). These ESKAPE and *E. coli* are major AMR pathogens in nosocomial settings.

A bacteriophage (phage) is a virus that infects bacteria. As soon as a phage adsorbs to the host’s cell wall, the phage genome is injected into the host cell. Temperate phages follow one of the two life cycles afterwards; the lysogenic cycle or the lytic cycle. In the lysogenic cycle, the phage genome is integrated into the host chromosome and it is called a prophage. In the lytic cycle, the prophage is induced to produce progeny phages in response to chemical or physical stressors (4, 5). Lysogenic phages or prophages are known to drive horizontal gene transfer (HGT) through transduction, but they also play an important role in increasing the genetic diversity of the host (6–10). Furthermore, defective prophages, also known as prophage-like elements, are stable in the host genome despite deleting most of the phage genes (6) and are known to increase host survival by conferring resistance against various stresses (11–13).

Plasmid conjugation has been well established as the major means of HGT of (AMR) genes (14), but recent studies have shed light on the role of phages in HGT of AMR genes since they are often encoded within the genome of the phage or prophage (15, 16). Metagenomic analysis revealed that AMR genes, such as ß-lactamases, are found in the phage genome (17–19). A clinical study reported that phages harboring AMR genes were identified in samples from patients with cystic fibrosis (20). Costa et al. reported that many prophage regions within the *A. baumannii* genome possessed several AMR and virulence factor (VF) genes using a bioinformatics approach (21). These studies have implied that phages and prophages probably transduce AMR genes more frequently than expected. However, the relationship between prophages and AMR genes has not been fully explored.

Pathogenic or VF genes in prophages have been mainly studied in *S. aureus* (22, 23), *E. coli* (24, 25), *Salmonella enterica* (26), and *Vibrio* spp. (27, 28), and their pathogenicity is associated with VFs encoded by the prophages. Other reports revealed that the expression of the Shiga toxin in *E. coli* (29) and mitogen factor in *Streptococcus canis* (30) depends on prophage induction. The studies mentioned above indicate that it is important to investigate the connection between host pathogenicity and their prophage to reveal their pathogenesis. However, there is very little known on the relationship between virulence and prophages in the rapidly emerging multidrug-resistant bacteria, such as ESKAPE pathogens and *E. coli*.

This study aims to understand the distribution of AMR and VF genes encoded in prophages, including the intact region, prophage-like elements, and satellite prophages (31) comprised in the bacterial genome, and discover their specific structural genomic features beyond the genera. We focus on seven clinically important AMR pathogens including ESKAPE pathogens and *E. coli* and analyzed their complete genomes deposited in a database to mine the prophage structure.

## Results

### Comparison of host genome size and the number of prophages

We investigated the correlation between the host genome size and the number of prophage elements present. The Pearson’s correlation coefficients (R-values) of each species were 0.57 for *A. baumannii*, 0.51 for *E. cloacae*, 0.76 for *E. coli*, 0.67 for *E. faecium*, 0.51 for *K. pneumoniae*, 0.67 for *P. aeruginosa*, and 0.63 for *S. aureus*. The R-values for all species were greater than 0.5, indicating that the number of prophages positively correlated with the host genome size (Fig. 1A).

**FIG 1.**
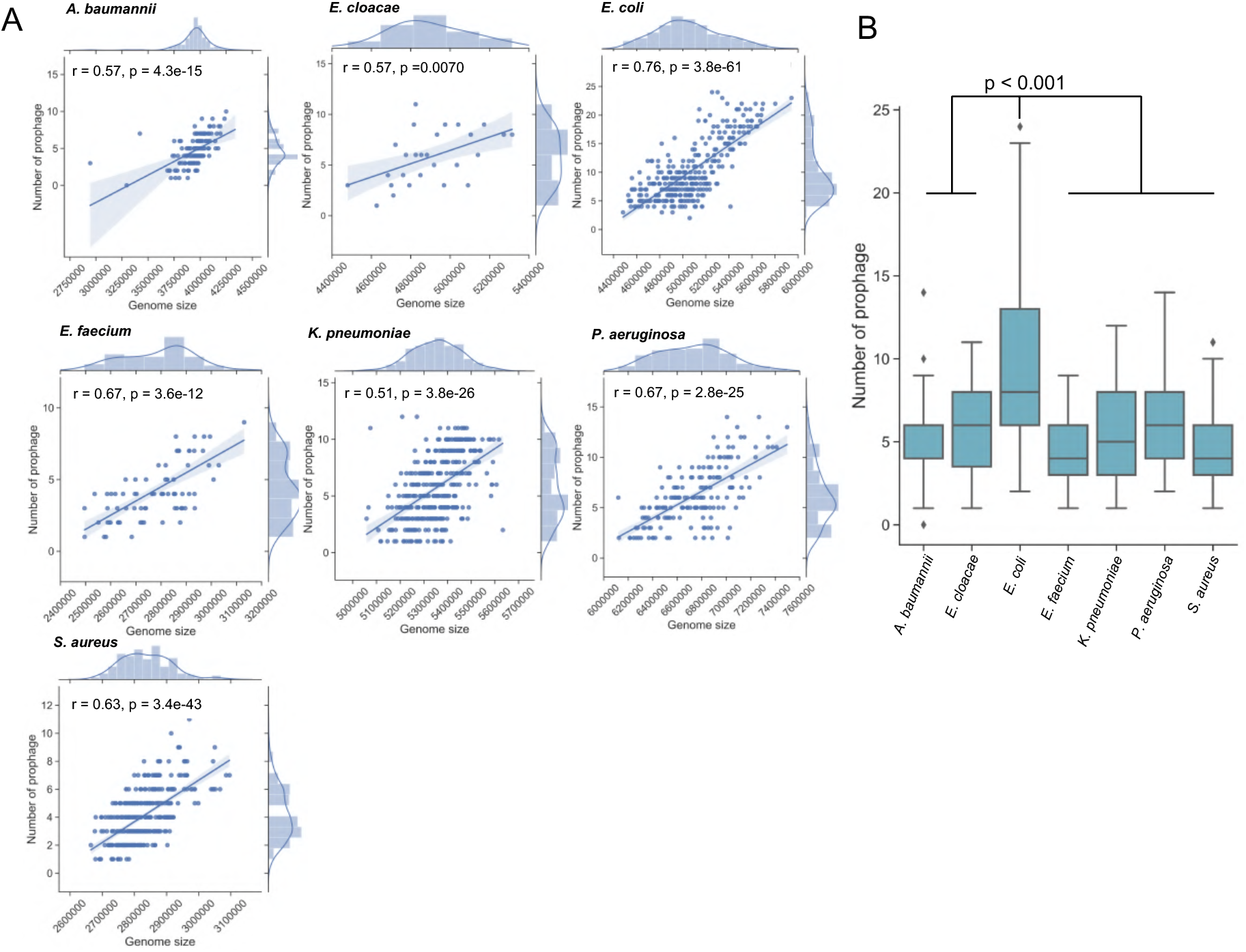
Comparison of genome length and the number of prophages. (A) Correlation and scatter plot of genome length excluding plasmids and the number of prophages. Plasmid samples were excluded to avoiding genome length bias. RefSeq and the complete genome were collected from the NCBI, and we evaluated the correlation between each genome length and the number of prophages obtained from PHASTER. r value indicates the Pearson’s correlation coefficient. (B) Box and whisker plot comparing the number of prophages for each species. The line inner box shows the median. Welch’s t-test was performed and P < 0.05 was considered as significant.

Next, we analyzed the number of prophages and prophage-like elements in each species. The number of prophages in *E. coli* was significantly higher compared to that in other species (Fig. 1B) (P < 0.001): *E. coli* possessed a maximum of 24 prophages (Accession numbers: CP027459 and CP024618) in which you can find prophages harboring Shiga toxin genes, *stx2A* and *stx2B*.

Screening for prophage elements in plasmids yielded a few cases, as depicted in Fig. S1. Comparison of the numbers of prophages in plasmids indicated that most *K. pneumoniae* plasmids harbored at least one or more prophages or prophage-like elements, and some *S. aureus* plasmids possessed one prophage or prophage-like element. However, *A. baumannii* possessed no plasmids harboring prophage or prophage-like elements. The number of prophages harbored in *K. pneumoniae* plasmids was significantly higher compared with that in *A. baumannii* and *S. aureus* plasmids (p = 4.7 = 10^-5^ and p = 0.0011, respectively) (Fig. S1). AMR genes from prophage elements were encoded in plasmids found in *E. faecium* (LC495616) and *K. pneumoniae* (AP018748, AP018752, AP018755 and AP018673), whereas no VF genes were detected on plasmid-harboring prophages in the accession numbers used in this experiment.

### The proportion of the number of AMR or VF genes to that of prophages in the host genome

We next examined the proportion of prophage elements encoding either AMR or VF genes. The proportions of prophages encoding VF genes were relatively high in *E. cloacae, E. coli, E. faecium*, and *S. aureus* (74.1, 65.4, 46.5, and 88.8%, respectively) (Fig. 2). In contrast, no prophages with VFs were detected in *A. baumannii*. Similarly, *K. pneumoniae* and *P. aeruginosa* also showed a low proportion of VFs harbored in corresponding prophages (0.8 and 11.6%, respectively) (Fig. 2).

**FIG 2.**
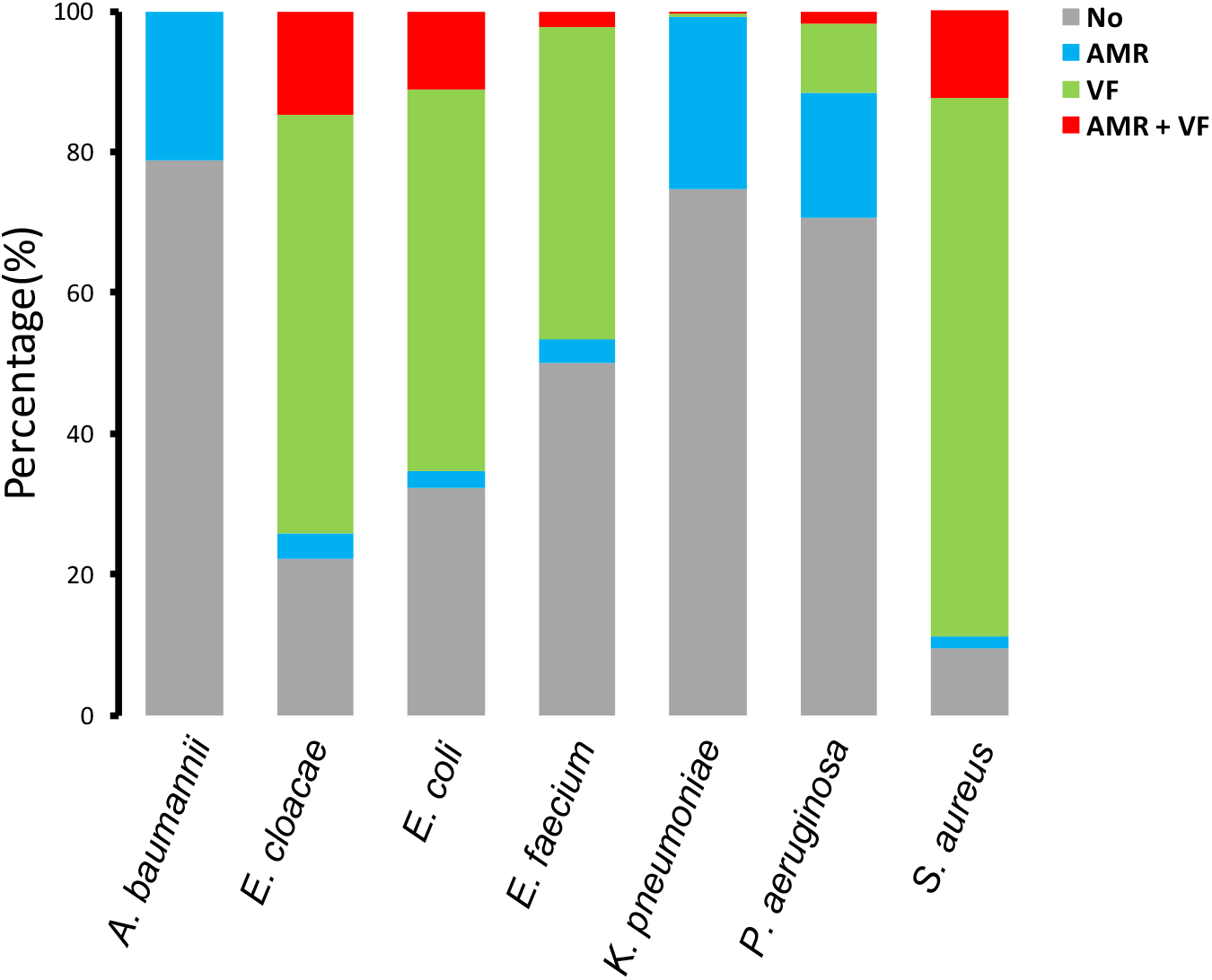
Proportion of the host genomes having prophages with AMR or VF genes. A bar chart showing the genome of each species that have the AMR and/or VF gene-containing prophages. Blue represents strains with AMR genes, green represents strains having VF genes, and red represents the strains having both AMR and VF genes. Gray shows strains having neither AMR nor VF genes.

Prophage regions encoding AMR genes were detected in all species. The proportions were 21.2% for *A. baumannii*, 18.5% for *E. cloacae*, 13.3% for *E. coli*, 5.8% for *E. faecium*, 24.7% for *K. pneumoniae*, 19.4% for *P. aeruginosa*, and 14% for *S. aureus*. The proportion of the host genome that contained both AMR and VF genes in prophages was 14.8% for *E. cloacae*, 11.1% for *E. coli*, 2.3% for *E. faecium*, 0.3% for *K. pneumoniae*, 1.7% for *P. aeruginosa*, and 12.3% for *S. aureus* (Fig. 2). Overall, the differences in proportions for each species indicated that the ratio between AMR and VF genes differed greatly depending on the species.

### Different prophage types encode either AMR or VF genes

We investigated the phage types integrated into the genome in each species. The name of each phage type was described using the most common phage from the PHASTER database. Phage types carrying AMR gene(s) are listed in Fig. 3 and aligned according to the number of phage types harboring AMR gene(s) (Fig. 3). The number of phage types harboring AMR genes was 46 (Fig. 3). Notably, the major phage types carrying AMR gene(s), Escher_RCS47 (RCS47), Entero_phi80, and Salmon_SJ46, were present in multiple species beyond the generic barrier. The Escher_RSC47 phage type was detected in all species except for *S. aureus*, and the Staphy_SPbeta_like phage type was detected in all species except for *E. cloacae* (Fig. 3).

**FIG 3.**
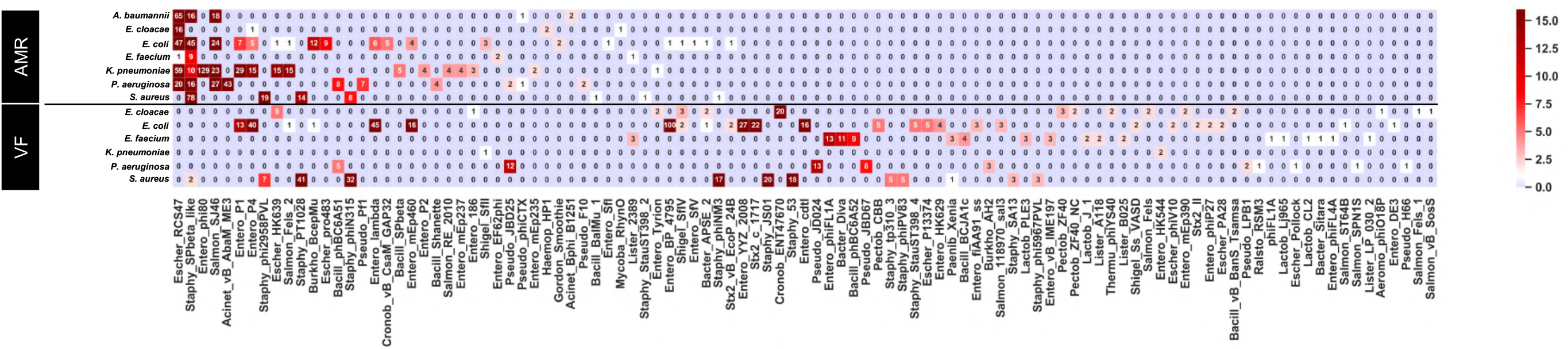
The type of prophages containing AMR and VF genes. Each prophage region that possessed AMR or VF genes was classified based on the most common phage in PHASTER. Heatmap shows the abundance of prophage type in each genus. The bottom represents prophage types described in PHASTER. The numeric character in each cell represents the number of detected phage names described in the most common phages on PHASTER. AMR, above the black line; VR, below the black line.

In contrast, we found that VF genes were widely distributed in a large number of phage types. There were 80 phage types, which were 1.7-fold higher than those harboring AMR genes. Furthermore, the prophage types carrying VF genes were different from those carrying AMR genes. Most of the phage types carrying VF genes, including ENT47670, phiFL1A, and Diva, were not detected in the prophages carrying AMR genes (Fig. 3). In addition, widely distributed VF-encoding prophages were not observed in various species. Overall, these results indicated that AMR and VF genes rarely coexisted within the same prophage and distribution pattern of prophage types containing AMR genes were different from that of VF-encoding prophage types.

### Comparing complete genomes of prophages harboring AMR and VF genes

We investigated and compared the completeness of prophage-encoding regions carrying AMR and VF genes. The regions were classified using criteria from PHASTER as follows; (1) intact, (2) questionable, or (3) incomplete. As shown in Fig. 4, percentages of prophages with AMR genes were 30.4%, 43.7%, and 27.1% for incomplete, intact, and questionable ones, respectively. Prophages carrying VF genes were mostly intact (70.3%), which indicated that the prophage retained most of its region, whereas 19.4 and 10.7% were incomplete and questionable phages, respectively (Fig. 4). In contrast to prophages carrying AMR genes, those carrying VF genes often encoded proteins that were crucial for the structure of the phage, such as the head, tail, and baseplate (Data Set S6 and S7). These phages were likely to produce progeny phages infecting other hosts. All prophages encoding Shiga toxins in *E. coli* were intact and contained phage structural proteins (Data Set S5). These data suggested that VF genes in prophages, such as the Shiga toxin gene could be easily transduced by a prophage.

**FIG 4.**
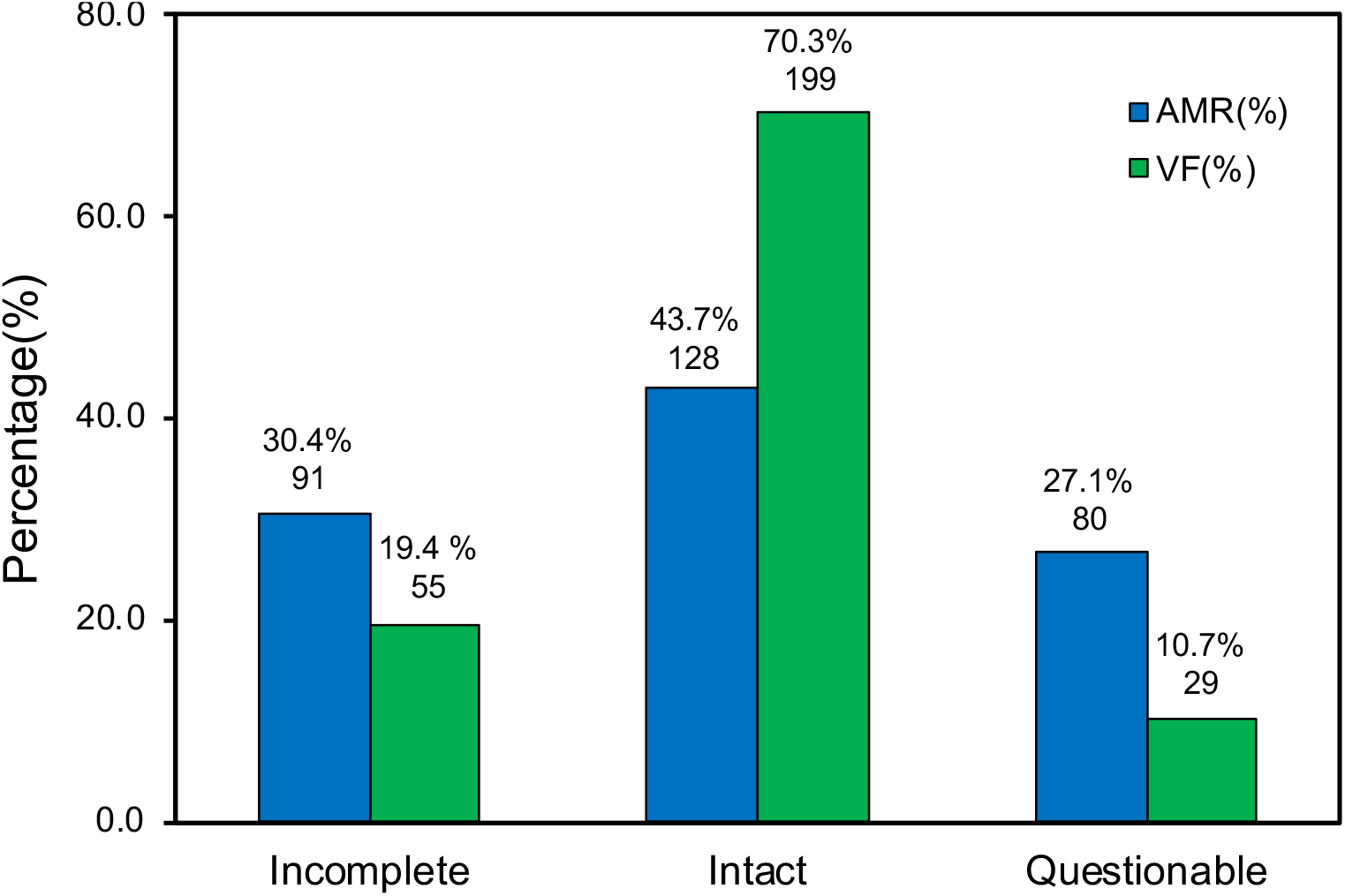
Analysis of prophage completeness using AMR or VF genes. Each prophage that had AMR or VF genes was classified as incomplete, intact, or questionable depending on the length and their completeness. The blue bar shows the percentage of AMR genes encoded in the prophage region, the green bar shows the percentage of prophages with a VF gene. Percentages regarded all AMR-or VF-encoding prophages as 100% and the numerical character below the percentage indicates the number of samples in each classification.

### Characterization of VF genes in prophages

To further investigate VF genes in prophage elements, VF genes for each accession number were examined using the VFDB from ABRicate and the presence of VF genes in prophages was visualized in the matrix (Fig. 5).

**FIG 5.**
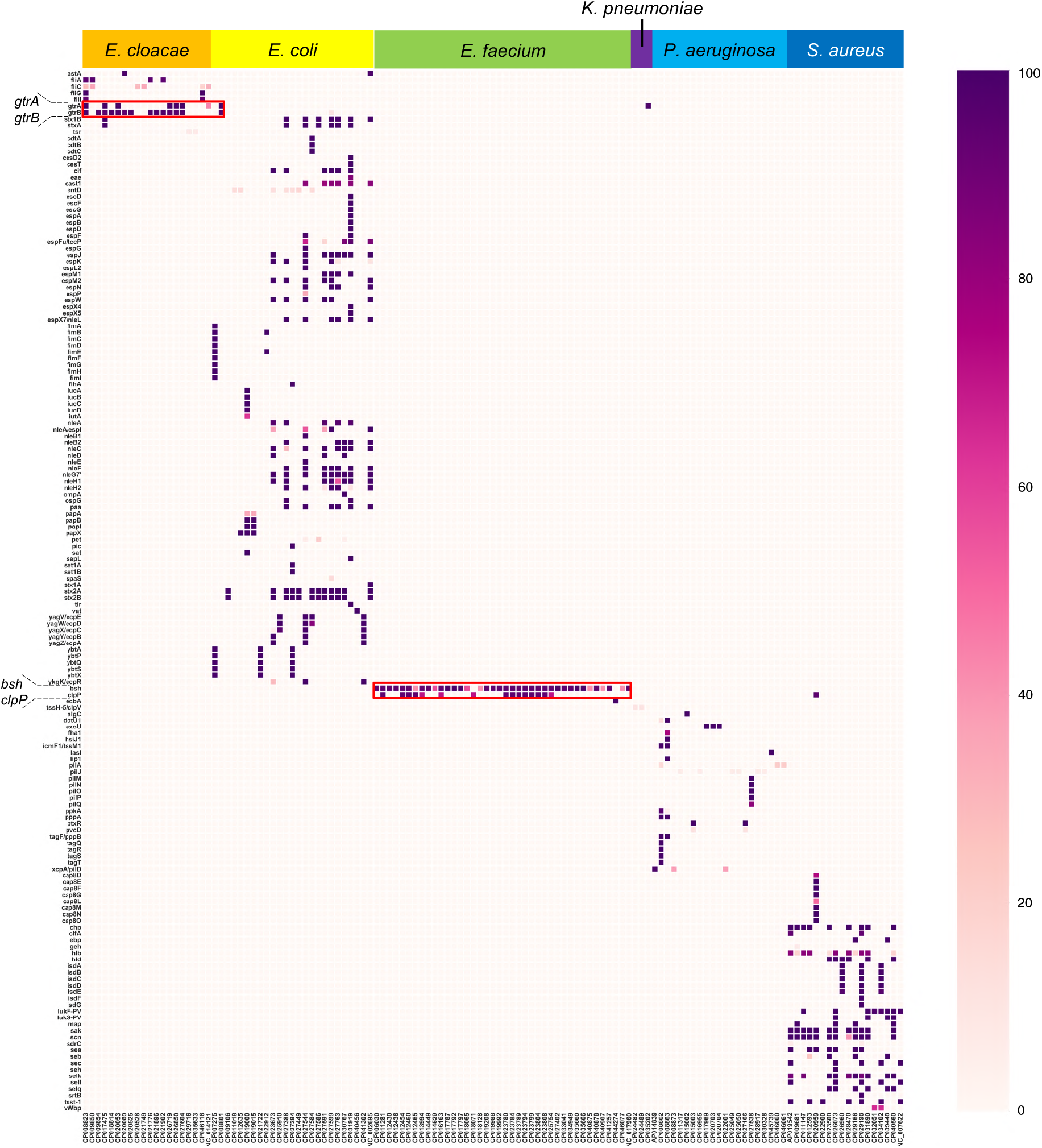
Distribution of VF genes in the prophage regions. DNA sequences of the prophage regions extracted from PHASTER were examined for the presence or absence of various VF genes using the VFDB database. The color density of the heatmap represents the coverage for each gene. The red frames show *gtrA, gtrB* and *bsh*. VF genes in *S. aureus* and *E. coli* are shown as the representative strains. All VF genes harbored by prophages are listed in Table S3.

Prevalence of prophage-encoded VF genes was unique to each species (Fig. 5). For example, we detected *gtrA* and *gtrB* in *E. cloacae* and *E. coli* strains and in just one *K. pneumoniae* strain but not in other species (Fig. 5). Bacteriophage ENT47670 (accession number: NC_019927) contained *gtrA* and *gtrB*. Interestingly, intact ENT47670 prophage, which also contained structural phage proteins, was detected in the prophage region where *gtrA* and *gtrB* were harbored (Data Set S5). This result suggested that *gtrA* and *gtrB* of the prophage region in *E. cloacae* were acquired through the transduction of ENT47670 or ENT47670-like phages (32) (Fig. 5; Data Set S5).

Most *E. faecium* had two specific genes in the prophage regions; *bsh* encoding a bile salt hydrolase and *clpP* encoding an ATP-dependent protease, but other species did not (Fig. 5). We mapped the prophage region harboring *bsh* and *clpP* in *E. faecium* using Easyfig (Fig. S2). We found that these genes were generally located at or near the gaps in the genome, and were scattered at various locations in the chromosome. These data suggested that coding regions *bsh* and/or *clpP* in prophages or prophage-like elements were successfully acquired via HGT. Classification of VF genes encoded by prophages revealed via VFDB keywords revealed that each species possesses functionally unique VF genes (Fig. S3).

### Characterization of AMR genes encoded by or nearby prophage elements

To investigate AMR gene distribution in the prophage elements of each strain in detail, AMR genes in the prophage region for each accession number were extracted using the ResFinder database from ABRicate and classified based on predicted substrates and mutation locations (See material and methods). Predicted substrates were described in Data Set S2. All species appeared to carry aminoglycoside modifying enzyme gene(s) and β-lactamase encoding gene(s) except for *E. faecium* (Fig. 6).

**FIG 6.**
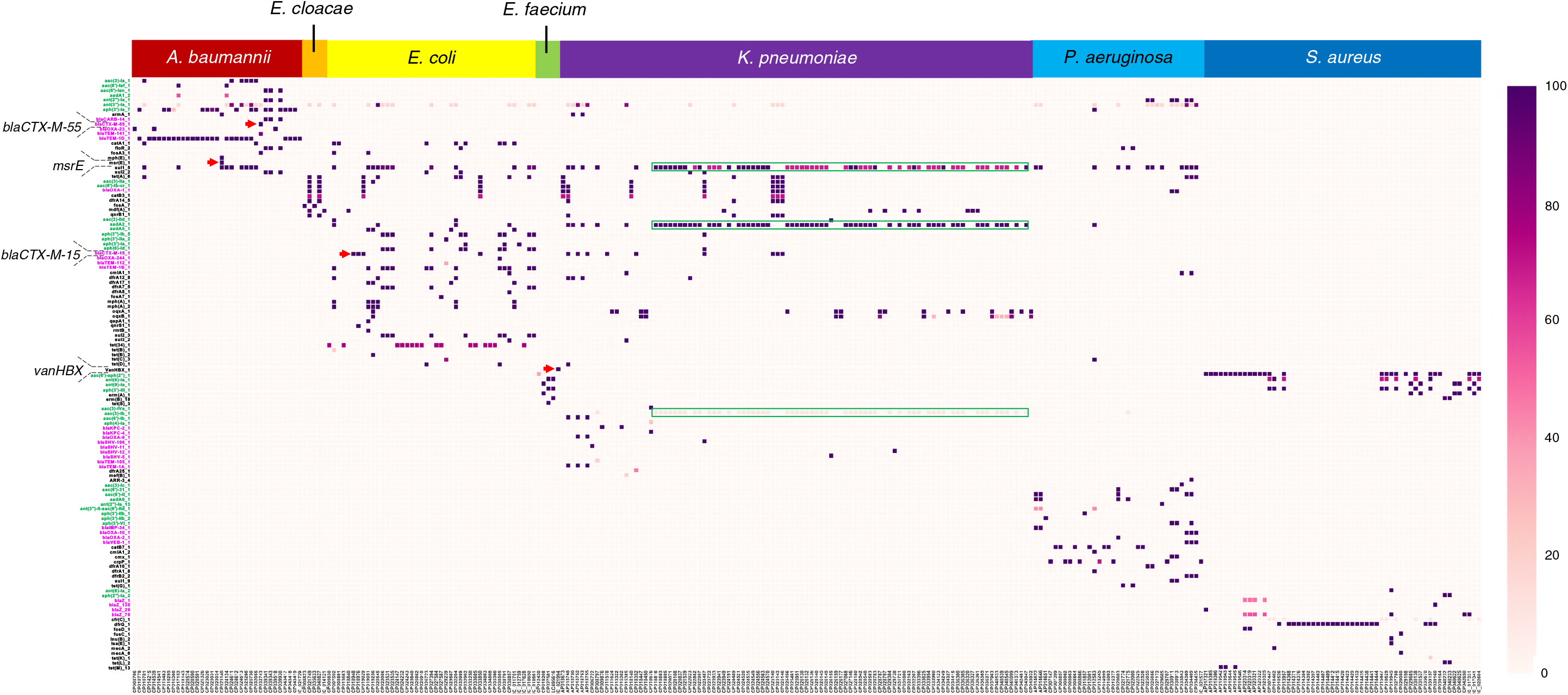
Distribution of AMR genes in the prophage region. We investigated the AMR genes in prophage regions extracted from PHASTER. AMR genes were detected using the ResFinder database. The color density of the heatmap represents the coverage of each gene. Green frame indicates the AMR genes cassette array detected in the *K. pneumoniae*. AMR genes, highlighted in green and magenta, show resistant genes against aminoglycoside and β-lactam, respectively. Magnified AMR gene names and red arrow represent AMR genes described in the results.

In contrast, species-specific AMR genes were also detected. For instance, *A. baumannii* harbored *msrE*, encoding the subunit for the ABC transporter and conferring resistance to erythromycin and streptomycin. *vanHBX*, conferring resistance to vancomycin, was detected in only *E. faecium*, and *vanHBX* was harbored in “intact” prophages with the genes encoding some structural proteins (Data Set S4). This data suggested that a part of *van* family of genes in vancomycin-resistant *E. faecium* were acquired via phage transduction. *bla*_CTX-M_ (33) was detected in the prophage region of several *K. pneumoniae* strains and only one *E. coli* strain (Fig. 6).

AMR gene cassettes were widely detected in various species. For instance, *K. pneumoniae* contained a cassette array of AMR genes including *sul1, aadA*, and *aacA* (Fig. 6). To detect the combination or the gene cassette of AMR genes harbored by prophage elements, we classified and clustered AMR gene cassettes for each prophage type (Fig. 7). Escher_RCS47, Salmon_SJ46, Acineto_vB_AbaM_ME3, and Staphy_SPbeta_like harbored various AMR genes (Fig. 7). In contrast, regions from prophages Entero_phi80, Entero_P1, and Escher_HK639 harbored a set of AMR genes including *sul1_5, aac (3)_lb_1*, and *aadA2_1* (Fig. 7). Another cassette array containing *ant(3”)-la_1, aph(3”)-la_7*, and *blaTEM-1D_1* was detected in Escher_RCS47, Staphy_SPbeta_like, and Salmon_SJ46. Since these AMR gene-combinations detected on prophage regions resemble the integron cassette array, we tried to identify the integrons in these prophage regions using the INTEGRALL database. We found that prophage regions in all species, except for *E. faecium* and *S. aureus*, possessed class 1 integrase. Therefore, we considered these characteristic regions containing AMR genes cassette array as an integron cassette array (Data Set S4).

**FIG 7.**
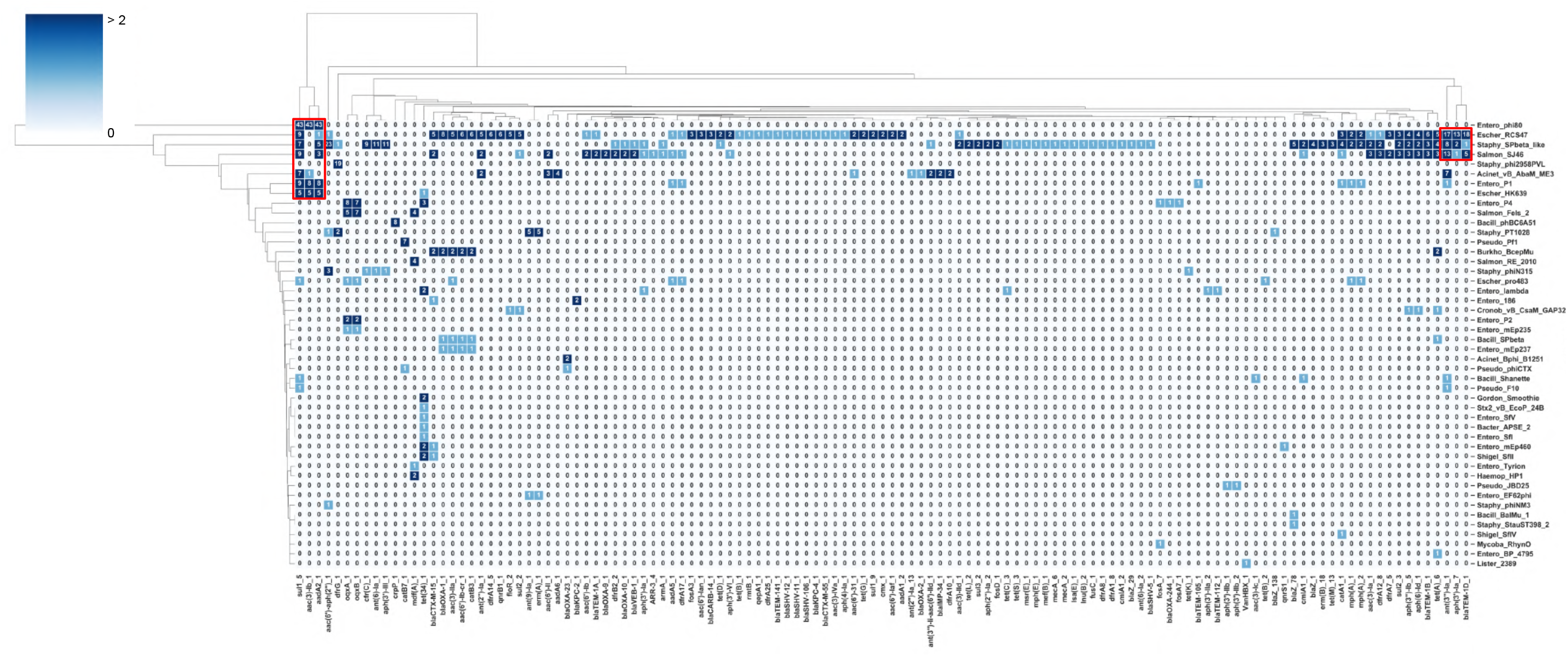
Cluster analysis of AMR gene-encoding prophages. Each prophage type included in all bacterial species used in this study was clustered under the same prophage name and visualized via a heatmap. The figure was created using seaborn, which is a Python module, and cluster method, which is a hierarchical clustering method (single linkage method). The numerical characters in cells mean the number of AMR genes in the indicated prophage. The red frames indicate the examples of cassette arrays of antibiotic resistance genes in the prophage region.

Next, we wondered if the prophage region containing integrons had a higher number of AMR genes than regions without integrons (Fig. 8). We examined the number of AMR genes in phage regions carrying integrons (Int/P, red), in those without integrons (No-int, green), and in strains where integrons were present somewhere other than the prophage region (No-Int/P, blue).

**FIG 8.**
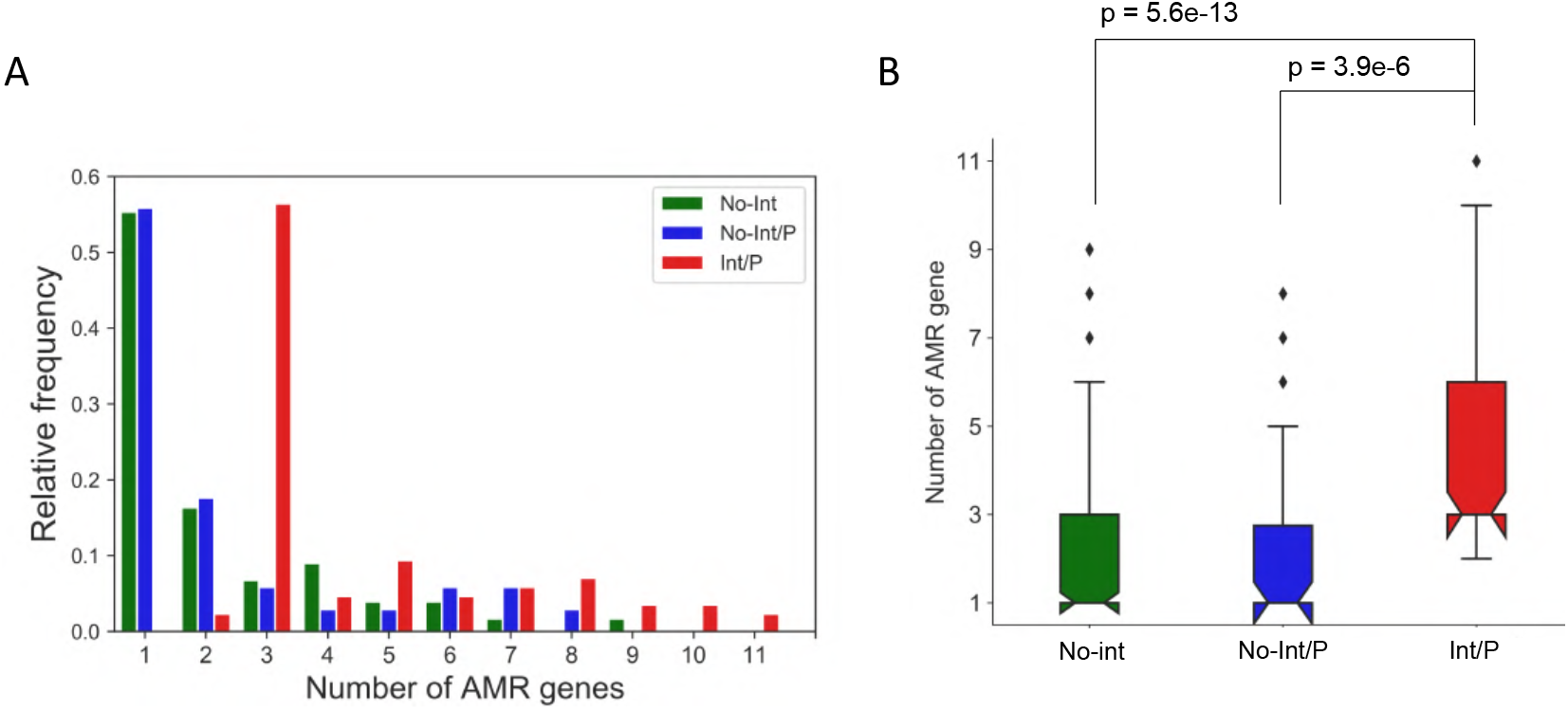
Comparison of the number of AMR genes with and without an integron. No-Int is a strain that does not have an integron, No-Int/P represents that the integron is located out of the prophage region, Int/P is a strain that has an integron in the prophage. Integron within a prophage was classified according to whether it has class 1 integrase or not. (A) The histogram shows the relative frequency of each number of AMR genes in each classification. (B) The boxplot shows the number of AMR genes in each classification. The notch of the boxplot represents the median, and Welch’s t-test was performed with P < 0.05 considered as significant.

Most of the phage regions (97%) carrying an integron possessed three or more AMR genes, whereas nearly 55% of phage regions without an integron possessed fewer AMR genes (Fig. 8A). These results indicated that the number of AMR genes were significantly higher in prophages with integrons than that in other groups (Fig. 8B).

### Structural features of prophage elements containing AMR genes

We briefly overviewed structural features of the prophage region and its association with AMR gene(s) by showing representative prophage sequences (Fig. 9). The prophage regions within the attachment sites (*attL* and *attR*) are shown in Figure 9. AMR genes in intact prophage regions existed either between or near integrase and/or transposase (Fig. 9A and B). Furthermore, AMR genes were located at the end of the prophage region, whereas the central position in the prophage region often encoded essential phage genes, especially structural proteins. Salmon_Fels_2, intact phage detected using PHASTER, lacked class 1 integron-integrase but phage-derived integrase was present just next to AMR gene(s) (Fig. 9A). We next visualized the prophage regions from Acineto_vB_AbaM_ME3, which were designated as incomplete or questionable phages using PHASTER and were termed as defective prophages (Fig. 9B). The prophage region from Acineto_vB_AbaM_ME3 had two different gene cassettes containing class 1 integrase and transposase, and both were located at the end of the region. The Acineto_vB_AbaM_ME3 type prophage region in strain CP020603 carried 11 AMR genes, which was the highest number per prophage-like elements in this study (Fig. 9B; Data Set S6).

**FIG 9.**
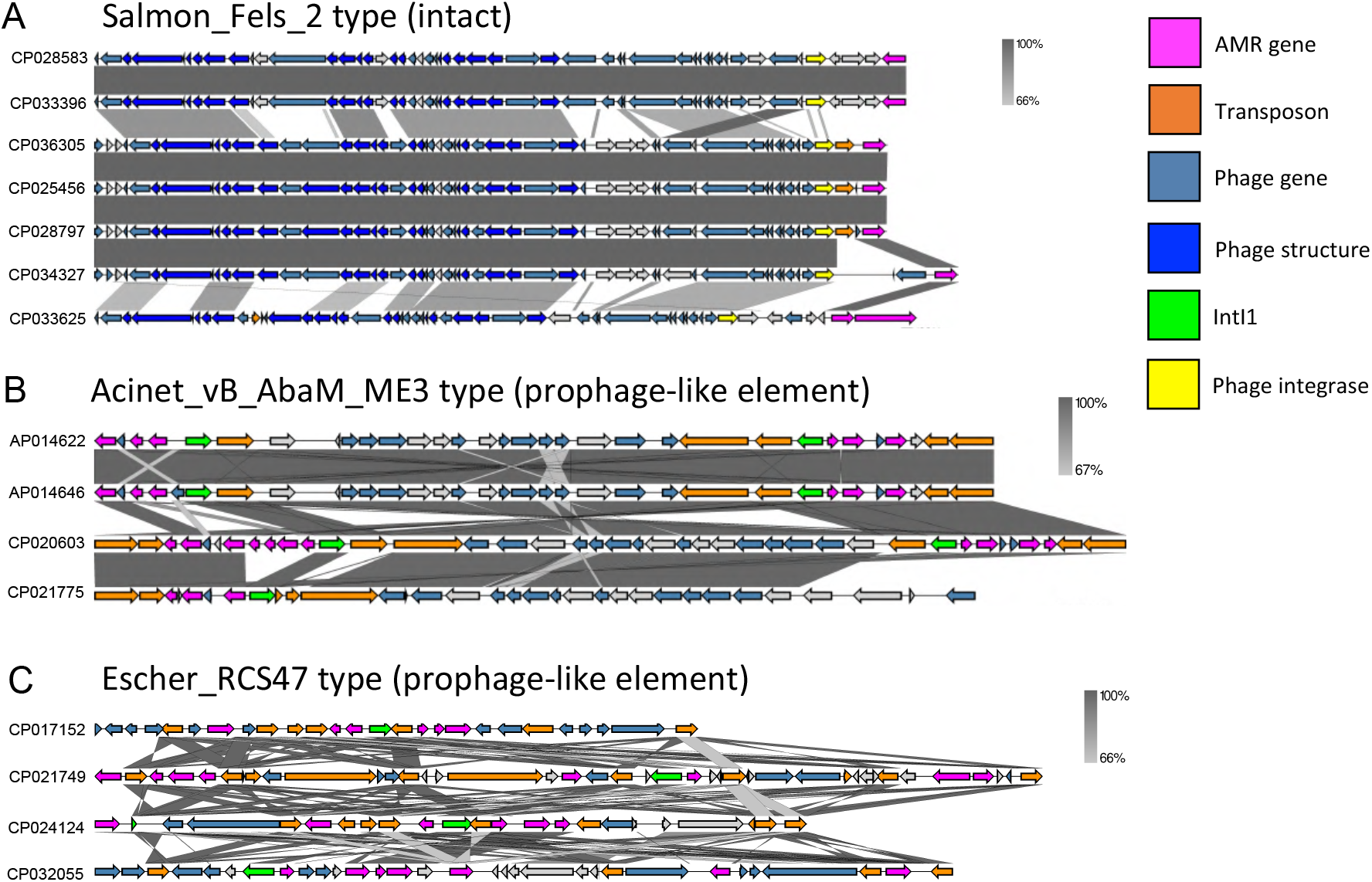
Comparative analysis of sequences and positional characteristics of AMR genes encoded by prophages. Representative prophage type names were selected based on genome completeness. Prophage sequences were defined via the attachment sites (*attL, attR*). (a) Salmon_Fels_2 prophage type was selected as an intact prophage; (b) Acinet_vB_AbaM_ME3 and (c) Escher_RCS47 were selected as prophage-like element (incomplete and questionable) prophages. The sequences of each prophage were selected at random and gray-shaded region represents the sequence similarity. IntI1 shows the class 1 integrase, which is shown by the green arrow, and phage-derived integrase is shown by the yellow arrow. AMR genes are shown in magenta. The figures were drawn using Easyfig 2.2.2.

In contrast to phage type described above, representative Escher_RCS47, which were classified as either incomplete or questionable, were devoid of most phage genes and their phage-derived transposases, and integrases were randomly arranged. Furthermore, AMR genes were located not only at the end but also at various positions including the center of the prophage sequence (Fig. 9C). The Escher_RCS47 prophage region was conserved in various species, and the characteristics of this prophage region might imply that these prophages had been integrated into the host genome for a long time. These results suggested that even though many phage structural genes had been deleted in evolutionary processes, the important genes for host survival, such as AMR genes, might have accumulated in such a conservative prophage region.

## Discussion

In this study, we collected the prophage elements from a total of 1,623 complete genomes of nosocomial AMR pathogens deposited in a public database and characterized AMR and VF genes harbored. Due to multi-drug resistance, ESKAPE bacteria have become a threat to global health, especially in elder patients. HGT agents, such as plasmids, prophages, and transposons often carry AMR and VF genes and move from bacteria to bacteria. Phage or prophage-carrying AMR genes are reported to be minor HGT agents (34, 35). In recent years, however, several studies have reported that AMR genes are present in the phage or prophage region (36–39), implying that they are associated with the prophage region more than expected. Nevertheless, there have been few reports comprehensively identifying and analyzing AMR/VF genes encoded in the prophage region of nosocomial AMR pathogens.

A previous representative metagenomic study has reported that phage regions encode almost no AMR genes (34). However, our study uncovered that the proportion of bacteria carrying AMR genes on prophage-like elements reached 10–20% in each species (Fig. 2). Our results suggested that the proportion of AMR genes encoded by prophages and defective prophages was higher than that of those encoded in the genome of phage particles and that AMR genes were preferentially accumulated in the prophage sequence and were stably inherited in the host genome (6). To date, plasmids are the most well studied HGT agent. Nonetheless, herein we did not compare the AMR gene richness between plasmids and prophage regions. Several prophage-like elements encoding AMR genes were inserted, and shared AMR genes in a plasmid; thus, we solely focused on the prevalence of AMR genes encoded by prophage regions, to shed light on previously overlooked contribution of prophages in AMR gene propagation.

We found that 57.5% of the prophages encoding AMR genes were defective prophages (defined as incomplete or questionable using PHASTER) (Fig. 4), and they lacked phage structural proteins (Fig. 4). Even if the prophage regions with AMR genes were determined as “intact”, these prophages often carried no structural genes (Data Set S4). One may speculate that these defective prophages cannot produce progeny phages due to the loss of structural proteins. However, Hayashi et al. has reported that several defective prophages encoding Shiga toxins in the genome of Enterohemorrhagic *E. coli* (EHEC) O157:H7 can produce progeny phages and that progeny phages can infect to other hosts (40). They have considered that prophages can complement each gene required for progeny phage propagation by sharing and using genes present in several prophage regions. Based on Hayashi’s report, we speculated that several defective prophages carrying AMR genes can complement the genes from each prophage region to synthesize progeny phages.

Prophage induction is triggered mainly by DNA damage via SOS response, and the host is lysed by the phage lysin. We found that more than 57.5% of AMR gene-harboring prophages were defective, whereas intact prophage was dominant in VF gene-harboring prophages (70.4%) (Fig. 4). A previous study has shown that prophages confer antibiotic resistance to the host and result in an increase of host viability (41). In other words, the host takes the advantage of AMR gene products of prophages for survival. A recent study has reported that the induction frequency of prophage carrying AMR genes decreases under the presence of antibiotics (42). As a result, prophages tend to become defective more frequently and are inherited in the host genome (43). In contrast, it has been reported that VF genes encoded in prophage can contribute to increasing phage infectivity by increasing the burst size and the latent period (44). In addition, prophages often harbor the genes involved in superinfection exclusion, which is a phenomenon that phage or prophage prevents infection by other phages (45, 46), and such genes are associated with host virulence (44). Therefore, we speculated that VF genes encoded in prophages have presumably more benefits for the prophage rather than bacteria, and the selective pressure of becoming defective is hardly caused in VF gene-encoding prophage. As a result, prophages containing VF genes tend to remain “intact” with phage structural genes.

Our results showed that most VF genes on prophage element were species-specific (Fig. 5), presumably because the mechanism of virulence to the host, such as the entry into the mammalian body (cell), and the toxicity differs depending on the species. Thus, each VF gene, which is responsible for virulence, could be detected in a species-specific manner. In contrast, modes of action of antimicrobials are common among bacterial species, and AMR genes are mobilized among bacterial genera through horizontal transfer; thus, similar AMR genes encoded in prophage elements are conserved in various bacterial species.

It has been known that genes for aminoglycoside modification enzyme and β-lactamase are widely conserved in various species (47, 48). In this study, these resistance genes were detected in or nearby prophages and prophage-like elements of almost all species, in accordance with previous studied (47, 48) (Fig. 6), suggesting that phages were related to HGT of these highly conserved AMR genes. A previous study showed that different subclasses of aminoglycoside nucleotidyltransferase (ANT) were detected in Gram-positive and -negative bacteria (49). While Gram-positive bacteria often possess ANT (4’, 6’, 9’), Gram-negative bacteria often have ANT (2”, 3”). In this study, ANT (6, 9) were detected on the prophage of *E. faecium* and *S. aureus*, but were rarely detected in Gram-negative bacteria (Fig. 6) in agreement with previous reports (49).

Duplication of the same AMR genes was detected in a single prophage region in particular, two or more aminoglycoside resistance genes with different mutations or subtypes were detected in a single prophage (Data Set S4). Duplication of AMR genes in prophage can increase the expression level and resistance to antibiotics (50). Furthermore, gene duplication or the increase of copy number is suggested to result in tolerance to mutations (51), accelerating evolution and genetic diversity (52). Thus, prophage element region may be a factory of producing AMR genes, which may result in the host rapidly becoming resistant to novel antibiotics.

We found that many prophages harboring AMR genes possessed an integron structure, and the number of AMR genes in integron-harboring prophages were significantly higher than that in prophage regions without an integron (Fig. 8). Some prophages with an integron have a characteristic structure with two different integron cassettes and a maximum of 11 AMR genes integrated into one prophage (Fig. 9). We proposed that the prophage region had a role as a reservoir for AMR genes. Integron structure was not detected in *E. faecium* and *S. aureus* prophage regions, and prophages without an integron structure often possessed a prophage-derived transposase near AMR gene(s). Interestingly, prophages RCS47, SJ46, and SPbeta-like phages, which have been detected in various species, harbor AMR genes and HGT-related genes in the original phage genome (53). Probably, phages carrying HGT-related genes could be key players to promote the transfer and accumulation of AMR genes. Even if the prophage region loses its infectivity due to genome incompleteness, AMR genes can be transferred by transposase (and integrase) derived from the phage. The Mu-like phage is present on a plasmid and the transposon Tn21 encoded in the phage region of the plasmid shared AMR genes (54). Overall, transposase derived from prophage has the potential to transfer AMR genes.

This study has some limitations. Some samples were collected from identical places, such as the same hospital or area, and the prophage regions of the genome from the same location were similar. Thus, further studies should focus on the relationships between the location and features of prophages. The sample number is uneven between each species. For example, the sample number of *E. cloacae* was 27, while that of *S. aureus* was 408. Since most of the complete genomes of *E. coli* are the EHEC, Stx (classified as “Toxin” in Fig. S3) and type III secretion systems were detected using the VFDB database, in agreement with previous reports (55, 56). Therefore, the distribution of VF genes and the percentage of Toxin and type III system harbored by prophages may have been influenced by the *E. coli* EHEC data. To analyze detailed and precise prophage sequence structure, we utilized a highly accurate prophage sequence using the National Center for Biotechnology Information (NCBI) RefSeq (57). We believe that our finding was endowed using not miscellaneous sequences but RefSeq, which curated highly accurate sequences.

Overall, we comprehensively detected AMR and VF genes in a wide range of strains, and our results will shed light on the important roles of phages as reservoirs and factors that transfer AMR/VF genes. Further research is needed to elucidate the amount of AMR and VF genes that are transferred to other strains via transduction in a clinical and natural environment.

## Materials and methods

### Data collection and prophage region detection

We compiled complete genomes and RefSeq data of seven bacterial species (169 sequences of *A. baumannii*, 27 sequences of *E. cloacae*, 324 sequences of *E. coli*, 88 sequences of *E. faecium*, 408 sequences of *K. pneumoniae*, 183 sequences of *P. aeruginosa*, and 424 sequences of *S. aureus*) from GenBank using Biopython version 1.76 (58). The genome size for each strain was referred to as the value described in GenBank (Data Set S1). Since *E. coli* had a large number of registrations on the NCBI, GenBank accession numbers were randomly selected so that the sample size would not increase. To detect prophages and prophage-like elements (28) for each strain, we used a custom application programming interface (API) from PHASTER (59). Prophage names were identified using the most common phage species described in PHASTER. Additionally, PHASTER was also used to classify prophages as either 1) “intact”, 2) “questionable”, or 3) “incomplete” based on the length of the phage region and the number of phage-derived genes (Data Set S4 and S5).

### Detection of AMR and VF genes

AMR and VF genes encoded by the prophage sequences were extracted using ABRicate version 1.0.1 (https://github.com/tseemann/abricate) under default settings. The ResFinder database (60) was used to detect AMR genes, and the Virulence Factors Database (VFDB) (61) was used to detect VF genes. VFs were classified based on VFDB keywords, and redundant (similar) keywords were summarized into short words (e.g., “adhesion,” “Apoptosis and Adherence,” and “Invasive” keywords were integrated into “Adherence,” and toxin-based genes were integrated into “toxin”). The same gene with a few, different mutations were distinguished as an accession number were assigned to each gene containing the respective mutation. Each of the genes was also assigned a “_ X” suffix.

### Integron analysis

Integrons and antibiotic cassette gene arrays in prophages were detected using INTEGRALL (62). Ambiguous integrases encoded in prophage region were examined using Blastp (protein-protein BLAST). If the amino acid identity was more than 80%, it was considered as a class 1 integrase. Numbers were assigned based on the order of the gene cassette and we referred to the number described in INTEGRALL. We classified the results into three groups, depending on the type of integron arrangement, as follows: 1) Prophage region with a class 1 integrase (Int/P); 2) prophage region without a class 1 integrase (No-Int); 3) integrase present, but did not exist in the prophage region (No-Int/P).

### Prophage elements and other data visualization

Prophage elements encoding *bsh* and *clpP*, were visualized using Blast Ring Image Generator (BRIG) version 0.95 (63). All BRIG parameters were set to default. Other prophage-related regions containing AMR genes were visualized using Easyfig version 2.2.2 (64). The thresholds of BLAST hits in Easyfig were analyzed using the default value. To visualize, we selected Escher_HK639, Entero_186, and Salmon_Fels_2, which were regarded as “intact” phages, encoding a phage structure protein. Alternatively, we selected Acineto_vB_AbaM_ME3, Entero_phi80, and Escher_RCS47, which were considered “incomplete” or “questionable” in PHASTER. The host accession number used for the analysis had three to five strains selected at random for each prophage. Other data were analyzed and visualized using Python 3.7.6 (Python Software Foundation, https://www.python.org), Matplotlib version 3.1.3, and Seaborn version 0.1.0.

### Statistical analysis

Pearson R correlation was calculated using the default jointplot function in seaborn. All statistical analyses were conducted using a two-sided Welch’s t-test using Python version 3.7.6 and SciPy Module version 1.4.1, and P < 0.05 was considered as a significant difference.

## Data availability

This study was performed using complete genomes registered on the NCBI. All accession numbers are listed in Data Set S1. All information on AMR and VF genes detected in this study are described in Data Set S2, S3. The information on prophages and AMR/VF gens including integron are summarized in Data Set S4 and S5. The presence of attachment sites (*attL, attR*) in each prophage region is described in Data Set S6, S7.

## Acknowledgments

We are grateful to Professor Yoshitoshi Ogura for his valuable suggestions on the manuscript.

## Supplementary material

**FIG S1.**
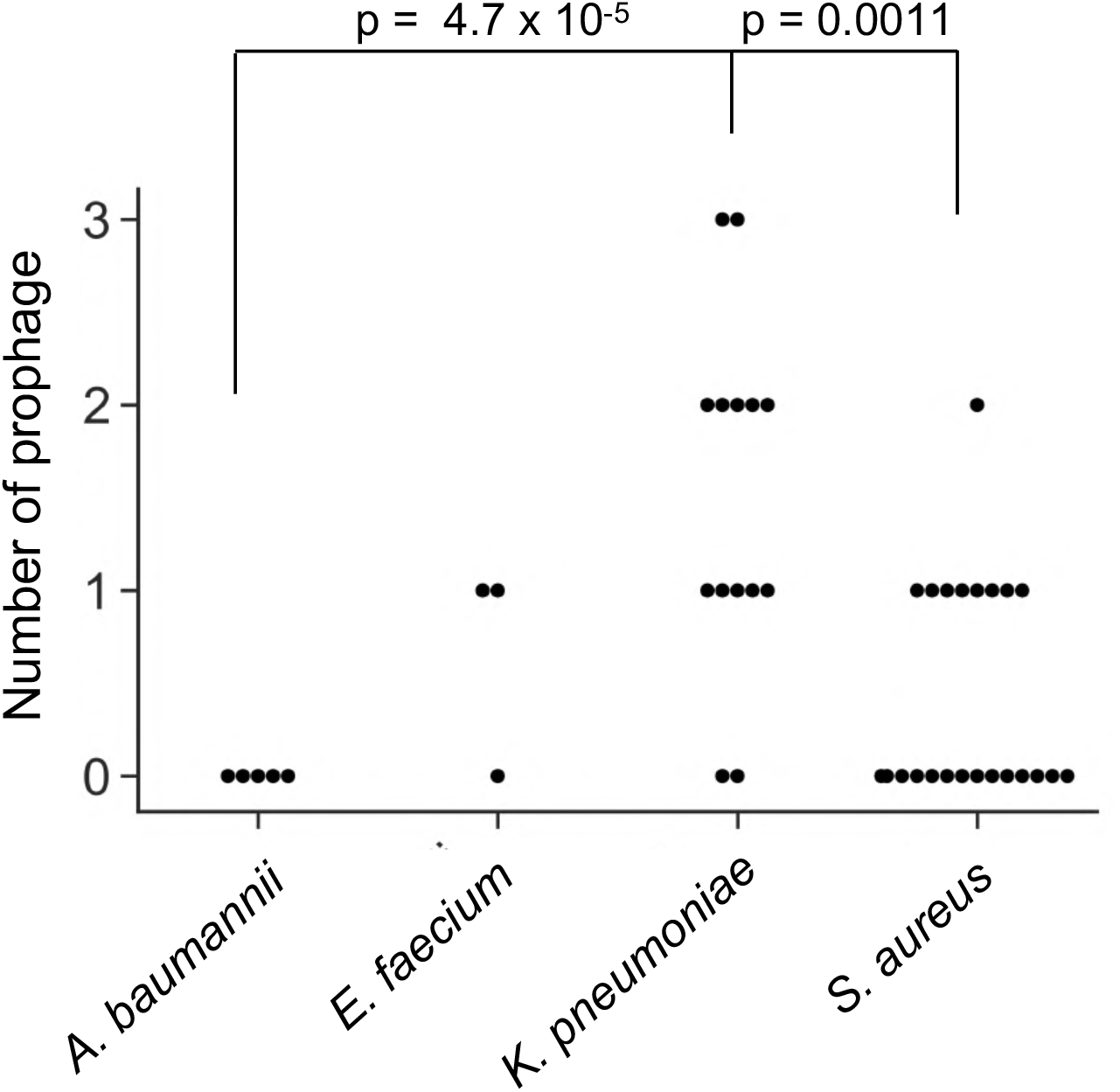
Comparison of the number of prophages present on the plasmid. The number of prophage regions on the plasmid was compared. Welch’s t-test was performed and P < 0.05 was considered significant.

**FIG S2.**
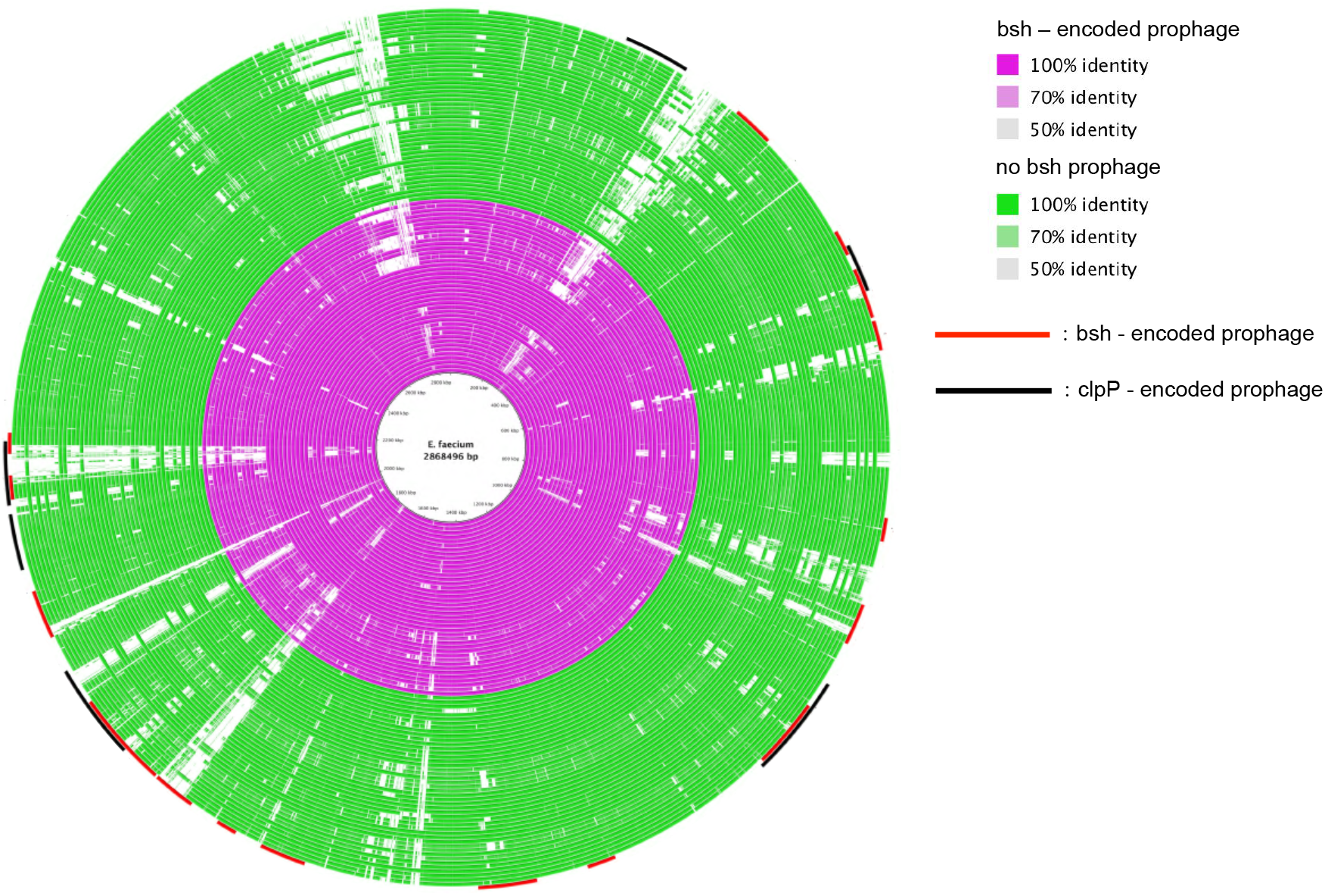
The mapping of prophages encoding *bsh* and *clpP* in *E. faecium*. The red circles (the inner circle) indicate the genome that includes *bsh*, and the green circles indicate the genome that does not include *bsh* in the prophage region. The red arc indicates the prophage region encoding *bsh*, and the black arc indicates that encoding *clpP*. The gap shows less homology (< 50 %) with the original genome. The figure was created using BRIG.

**FIG S3.**
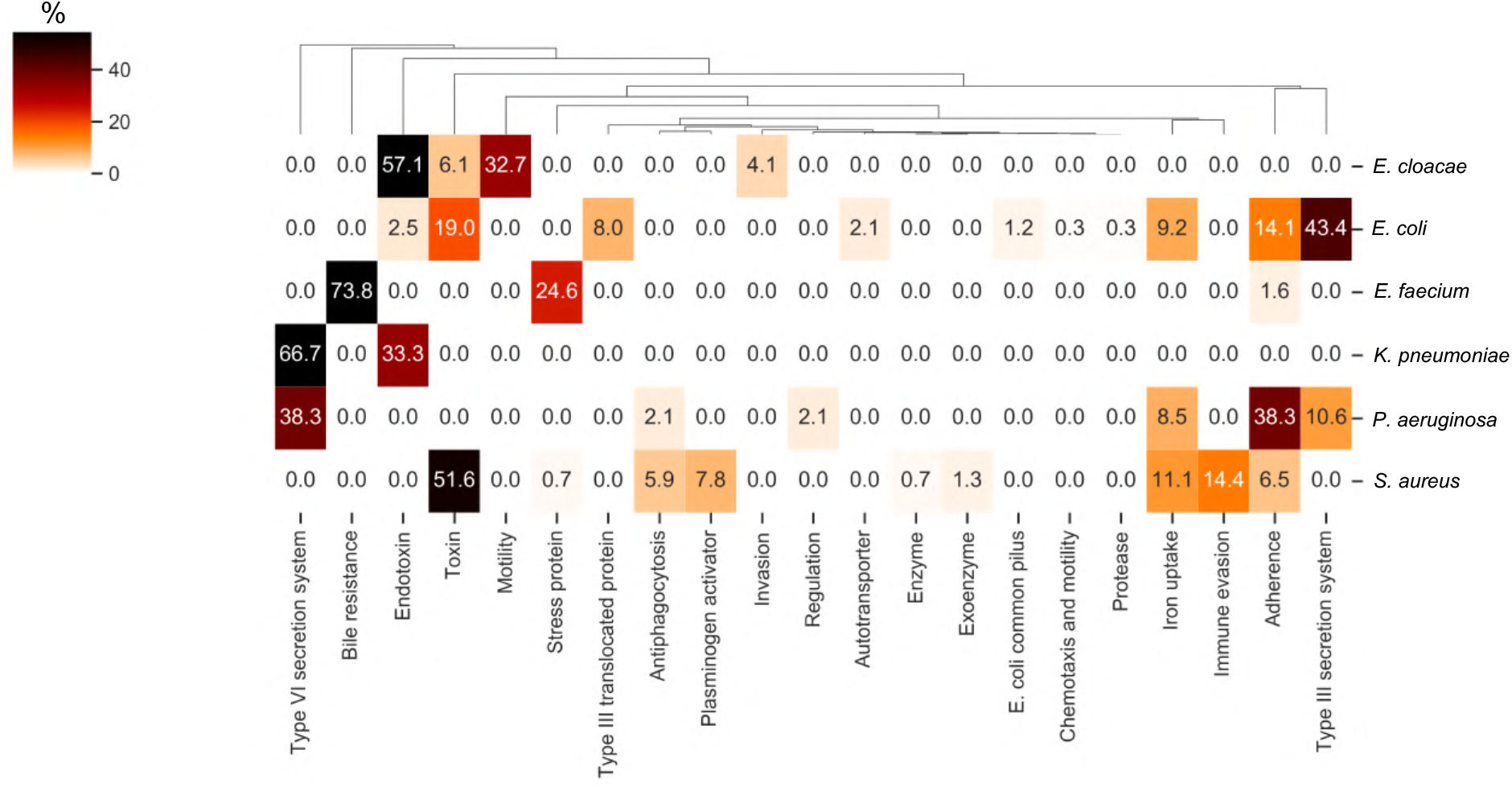
Classification of VF genes encoded by prophages. Each VF gene was classified based on the keywords described in the VFDB. VF genes harbored by prophages were not found in *A. baumannii*. The numerical value in the heat map indicates the percentage (%) classified into keywords for each strain.

